# Transplantation in neonate mouse recipients enhances umbilical cord blood CD34^+^ cells permissiveness to ETO2::GLIS2-driven transformation

**DOI:** 10.64898/2026.01.12.698953

**Authors:** Klaudia Galant, Vilma Barroca, Saiyirami Devanand, Didier Busso, Guillaume Piton, Nathalie Dechamps, Laurent Renou, Thomas Mercher, Stéphanie Gachet, Françoise Pflumio

## Abstract

Acute megakaryoblastic leukemia (AMKL) is a rare and aggressive pediatric acute myeloid leukemia subtype, frequently driven by the ETO2::GLIS2 (EG) gene fusion in de novo AMKL patients. The EG fusion confers poor prognosis and high relapse risk. In a previous study, we demonstrated that human fetal hematopoietic cells are intrinsically permissive to EG-driven transformation and that microenvironmental cues critically enhance leukemogenesis in less permissive human cord blood (CB) cells. However, most pediatric AML models are established in adult recipient mice, potentially overlooking developmental niche-specific effects. Here, we investigated whether mouse neonates provide a more permissive environment for EG-driven leukemogenesis from human CB hematopoietic stem and progenitor cells (HSPCs). Comparison of xenotransplants of EG-transduced CB CD34^+^ cells into sublethally irradiated NSG neonate or adult mice show that neonatal recipients are more supportive of EG-driven leukemia when cell doses are adjusted for recipient size. Neonates exhibited a higher frequency of leukemia-initiating cells (LIC), enhanced expansion of EG/GFP^+^ leukemic cells, and increased disease penetrance compared with adults at similar dose/size ratio, despite similar overall human hematopoietic engraftment. Leukemic cells arising in neonates and adults displayed comparable phenotypes and secondary transplantation capacities, indicating functional equivalence. Both CD34^+^CD38^-^ and CD34^+^CD38^+^ CB subpopulations were sensitive to EG-driven transformation, with neonates consistently showing higher LIC frequencies than adults. Together, our findings demonstrate that the developmental stage of the host microenvironment is a critical extrinsic determinant of EG-mediated leukemogenesis. Neonatal recipients constitute a physiologically relevant and highly sensitive model for studying pediatric AMKL initiation with low cell inputs, enabling refined investigation of microenvironmental signals that promote leukemia development and may reveal novel therapeutic vulnerabilities

## Main

Acute megakaryoblastic leukemia (AMKL) is a rare but aggressive acute myeloid leukemia (AML) subtype mostly found in children before the age of 3^1–3^. *De novo* AMKL patients carry chromosomic rearrangements, including the *ETO2::GLIS2* (EG) fusion in 25-30% of cases^4^. This fusion is associated with a poor prognosis and a high risk of relapse^4–6^. Both murine and human cell models have indicated that fetal cells are highly permissive to EG-driven transformation^7,8^. When using human cord blood CD34^+^ cells, specific human cytokines drastically enhanced the leukemogenicity^8^. Together, both the cell of origin and the microenvironment may contribute to the emergence of EG-driven leukemia across different developmental stages.

While pediatric AML are frequently modelled by injection of normal hematopoietic cells into adult recipient mice^7–15^, the recipients age may be of particular importance when studying the initiating steps of pediatric leukemogenesis. Indeed, fetal, newborn and adult hematopoietic cell microenvironments vary in terms of anatomical localization, niche cell composition and in their capacity to support hematopoiesis^16–18^. Therefore, newborn mice (age ≤ 3 days) might be interesting recipients for studying the development of pediatric leukemia, taking into account both intrinsic and extrinsic fetal specificities. Several studies have focused on the differences between neonate and adult recipient mice, encompassing both normal and pathological hematopoiesis^19–23^. Intrahepatic (IH) injections in neonates provides the possibility to mimic hematopoietic stem and progenitor cells (HSPCs) residency in the fetal liver and then their physiological migration to the bone marrow, thereby modelling the hematopoietic transition from fetal to adult niches. Moreover, studies have shown that immunodeficient mouse newborns exhibit better engraftment of human AML cells after sublethal irradiation than adult recipients, suggesting that they may provide an improvement for pediatric AML xenograft models^19,22^. Therefore, based on our recent results that umbilical cord blood (CB) cells greatly rely on microenvironmental cues to efficiently transform when expressing the EG gene fusion^8^, we tested here whether neonate recipients were more permissive to CB HSPC EG-driven leukemogenesis than adult recipients.

We reasoned that neonates are much smaller than adults (NSG mice, neonates: ∼2.5g two days after birth vs adults: 25-30g), thus injecting the same (large) number of cells into neonate mouse livers might result in a much stronger imbalance between the injected cells and organ cells. Therefore, CB EG-transduced CD34^+^ HSPCs were transplanted side-by-side into neonate or adult sublethally irradiated NSG recipients each with different doses. We used intrahepatic and intravenous routes for neonate and adult mice, respectively, with a common dose of 1x10^5^ EG/GFP^+^ cells per mouse in both neonates and adults. We also tested a 1x10^4^ EG/GFP^+^ cell dose in neonates, and, inversely, a 1x10^6^ GFP^+^ cell dose in adults (Figure 1A). A lethal disease developed in some (2/5 recipients, 40%) adult NSG mice receiving 1x10^5^ EG/GFP^+^ cells as previously shown^8^, whereas increasing ten times the number of EG/GFP^+^ cells led to fully penetrant disease in all (9/9, 100%) adult mice (Figure 1B). In neonates, 1x10^5^ EG/GFP^+^ cells induced a lethal pathology in 16/17 (94%) recipients, while 1x10^4^ EG/GFP^+^ cells induced leukemia in 3/13 mice (23%) (Figure 1B). Importantly, there was no difference in the survival of neonate vs adult mice if the cell dosage was adapted to the weight of the recipient (namely 1x10^5^ cells for neonates vs 1x10^6^ cells for adults or 1x10^4^ vs 1x10^5^ cells, respectively) (Figure 1B). As a result, the frequency of leukemia-initiating cells (LIC) was 6 times higher in neonate vs adult mice (Figure 1C). When comparing both neonate and adult recipients injected with the highest cell dose, respectively 1x10^5^ and 1x10^6^ GFP^+^ cells/mouse, equivalent robust percentages of hCD45^+^ cells were detected in the bone marrow (BM) at the time of disease development and sacrifice, whereas recipients injected with less cells (1x10^5^ and 1x10^4^ cells/mouse) had lower percentages (Figure 1D). We focused on neonate and adult recipients receiving the highest EG/GFP^+^ cell doses (1x10^5^ vs 1x10^6^ respectively, 22-55% GFP^+^ cells at the start, Figure 1A) to characterize the engrafted human cells. Percentages of EG/GFP^+^ cells were significantly higher in mice injected at the neonate stage (Figure 1E), suggesting that the young environment induced superior development of EG/GFP^+^ cells. hCD45^+^ EG/GFP^+^ cells exhibited the same high KIT expression and variable levels of CD34, CD56 and CD41 surface markers regardless of the recipient age/size (Figure 1F, Figure S1A-B), a phenotype that we previously defined as characteristics of transformed cells^8^. hCD45^+^ GFP^-^ cells included CD19^+^ B cells, CD3^+^ T and CD11b^+^ myeloid cells (Figure 1G,, Figure S1A-C), as typically observed for normal hematopoietic cells in humanized NSG models^24^. Thus, when injecting the same number of EG/GFP^+^ cells in neonate and adult mice, a more efficient leukemic cell generation was observed in neonates. Interestingly, and in line with the cell phenotype, these leukemic cells had similar secondary transplant abilities in adult recipients, independently of their original growing niche. Indeed, phenotypically similar abnormal cells isolated at sacrifice from originally injected neonate and adult recipients induced disease development at the same rate and kinetic (Figure 1H), indicating they were functionally equivalent transformed cells. Thus, taking the weight difference between adult and neonate mice in consideration results in similar human cell levels but higher abnormal cell development in newborns, supporting neonatal environment as being more advantageous to disease set-up. Of note, there was no difference between female and male neonate recipients in terms of survival (Figure S2A), hCD45^+^ and EG/GFP^+^ cell engraftment (Figure S2B-C) and phenotype (Figure S2D-E), in contrary to previous reports in adult mice^25,26^. Also, different injection routes (i.e. intra-femoral in adults vs intravenous in neonates) did not modify these observations (Figure S3A-F), suggesting they were not critical determinants of EG-driven CB transformation.

**Figure 1.**
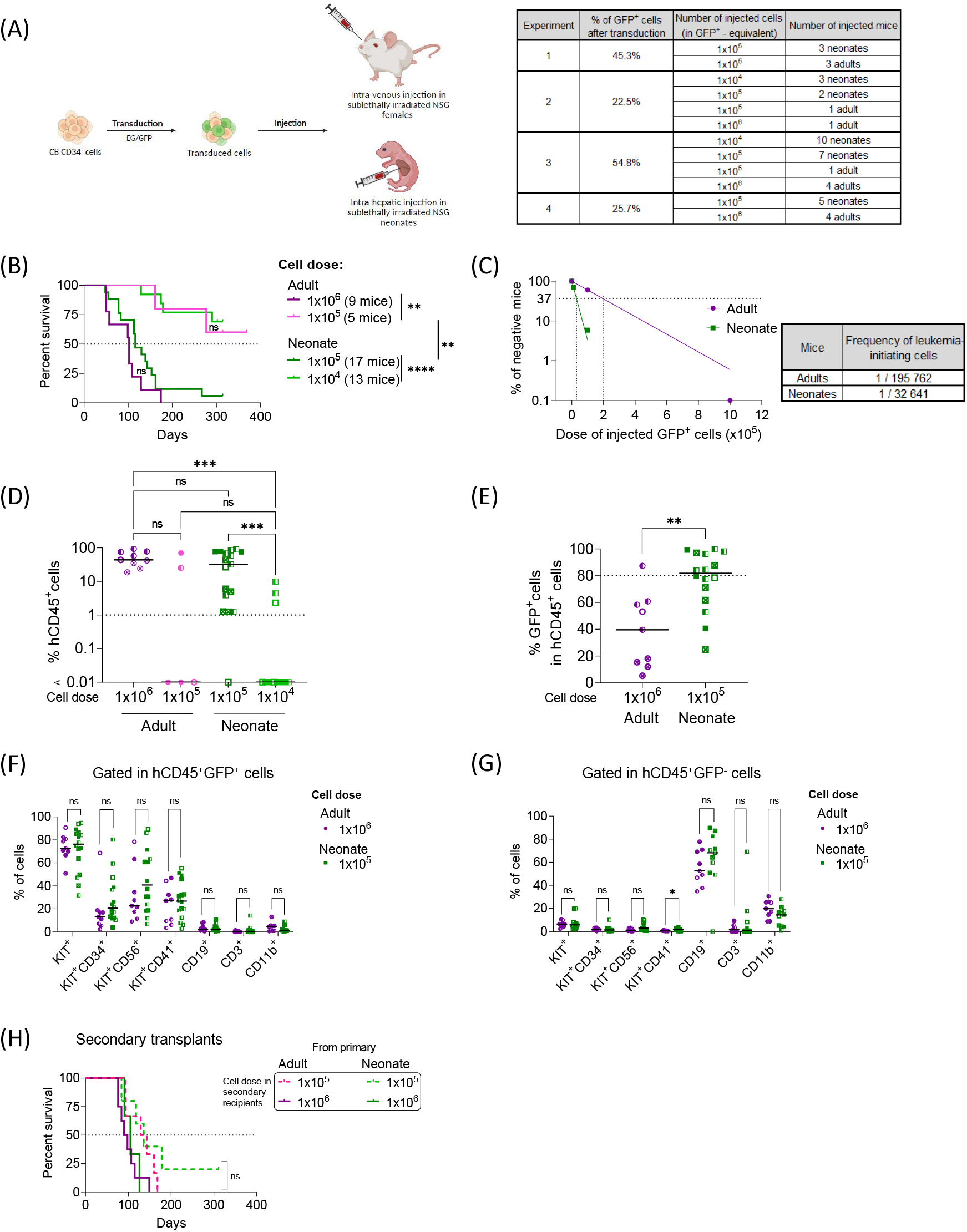
Increased EG-GFP^+^ cell numbers enhance the leukemogenic potential of CB EG cells *in vivo*. (A) Experimental design (*Created in BioRender. Gachet, S. (2026)* https://BioRender.com/2xx786y). CB CD34^+^ cells were transduced and injected into sublethally irradiated (1Gy) NSG neonates (48h after birth by intrahepatic route, females and males, 1x10^4^ or 1x10^5^ GFP^+^ cells per mouse) or sublethally irradiated (2Gy) NSG female adults (intravenous route, 1x10^5^ or 1x10^6^ GFP^+^ cells per mouse). Neonates were weaned at the age of 28 days. Mice were monitored daily and euthanized when presenting disease symptoms (hind limb paralysis, signs of anemia, loss of weight) (left panel). Table recapitulating the number of injected mice/condition. n=3 biological CB EG replicates in adult mice. n=4 biological replicates in 1x10^5^ GFP^+^ cell-injected neonates. n=2 biological replicates in 1x10^4^ GFP^+^ cell-injected neonates. The transduction efficiency (% of GFP^+^ cells after transduction) is indicated (right panel). (B) Kaplan-Meier survival plot of NSG mice injected with EG-transduced CB cells. Median survival was 102 days for adults receiving 1x10^6^ GFP^+^ cells, 116 days for neonates receiving 1x10^5^ GFP^+^ cells. (C) Frequency of leukemia-initiating cells in adults vs neonates. The fraction of non-responding (healthy) mice was plotted vs the number of injected CB EG-GFP^+^ cells. The frequency of leukemia-initiating cells was estimated using the L-Calc software (StemCell Technologies). (D) Percent of engrafted human CD45^+^ cells in the long bones BM of studied mice. (E) Percent of GFP^+^ cells in the long bones BM of studied mice (gated in hCD45^+^ cells). (F) Percent of KIT^+^, KIT^+^CD34^+^, KIT^+^CD56^+^, KIT^+^CD41^+^ and normal CD19^+^, CD3^+^ and CD11b^+^ cells in the long bones BM of deceased recipients (gated in hCD45^+^ GFP^+^ cells). (G) Percent of KIT^+^, KIT^+^CD34^+^, KIT^+^CD56^+^, KIT^+^CD41^+^ and normal CD19^+^, CD3^+^ and CD11b^+^ cells in the long bones BM of studied mice (gated in hCD45^+^ GFP^-^ cells). Symbols used in (D-G) show the different experiments depicted in (A). (H) Kaplan-Meier survival plot of secondary adult NSG mice injected with 1x10^5^ or 1x10^6^ hCD45^+^ GFP^+^ cells recovered from deceased primary mice (adults or neonates). Median survival was 136 days for secondary recipients receiving 1x10^5^ GFP^+^ cells from either adult or neonate primary deceased recipients, 94 days for secondary recipients receiving 1x10^6^ GFP^+^ cells from adult primary deceased recipients and 105 days for secondary recipients receiving 1x10^6^ GFP^+^ cells from neonate primary deceased recipients. Statistical significance is indicated as p values (B and H: Log-rank test, D: Kruskal-Wallis test, E-F-G: Mann-Whitney test). ns: not significant, **: p<0.01, ***: p<0.001.

Because of the low frequency of HSPCs permissive to EG transformation in CB (Figure 1C) and since previous works in mouse pointed the permissive cells to reside within the hematopoietic stem cells (HSC) compartment^7^, we tested EG-transduced flow-sorted CD34^+^CD38^-^ cells (HSC/MPP compartment, thereafter referred to as CD38^-^ cells) and CD34^+^CD38^+^ cells (progenitor compartment, thereafter referred to as CD38^+^ cells) for their ability to get transformed in neonate and adult NSG mice. We also tested different cell doses to account for the recipient size ratio (Figure 2A-B). Thus, adult mice received an equivalent of 1x10^5^ and 1x10^4^ EG/GFP^+^ cells, and neonates were given 1x10^4^ and 1x10^3^ EG/GFP^+^ cells of CD38^-^ /CD38^+^ sorted cells (Figure 2B). Interestingly, the survival of neonates and adult mice injected with EG/GFP^+^CD38^-^ cells closely mimicked that of mice injected with bulk EG/GFP^+^CD34^+^ cells (Figure 2C, Figure 1B). Likewise, the number of injected cells was important for CB EG-driven leukemogenesis. Indeed, injecting low (1x10^4^ in adults and 1x10^3^ in neonates) doses of transduced cells was less efficient in promoting disease whereas 10-fold higher cells (1x10^5^ in adults and 1x10^4^ in neonates) induced a fully penetrant disease (Figure 2C, Figure S4A-D). Besides, 100% of adults given 1x10^5^ EG/GFP^+^CD38^+^ cells developed a disease with full penetrance (4/4 mice), compared with 57% (4/7) pups injected with 1x10^4^ EG/GFP^+^ cells (Figure 2D), indicating that CB CD38^+^ progenitor cells also support EG leukemogenesis when injected at the right doses, although the adult environment may be more permissive for CD38^+^ derived cells. Injecting 10-times less cells into adults or neonates resulted in a similar disease latency (Figure 2D, Figure S4A-D). When calculating LIC frequency, no difference was observed between CD38^-^ and CD38^+^ derived cells, still neonates had higher frequency over adult mice (Figure 2E-G). Measurement of hCD45^+^ cell percentages (Figure 2H-I), GFP^+^ cell proportion (Figure 2J-K) or phenotype of hCD45^+^ EG/GFP^+^ abnormal or hCD45^+^ GFP^-^ normal cells (Figure S4E-F) did not reveal significant differences between neonates and adult recipients at time of disease development and sacrifice. However, there was a trend to enhanced EG/GFP^+^ cell percentages in neonates with higher mouse number with high percentage of EG/GFP^+^ cells (Figure 2J-K) that can be explained by the superior support of the neonate environment. These data suggest that both CB-derived CD38^-^ and CD38^+^ fractions are sensitive to EG transformation. As these CD38^-^ and CD38^+^ cell populations can be subdivided in HSC, MPP, CMP, and progenitors^27,28^, testing the sensitivity of even more purified progenitors would be required to assess the EG-driven leukemia cell of origin potential of specific human subpopulations.

**Figure 2.**
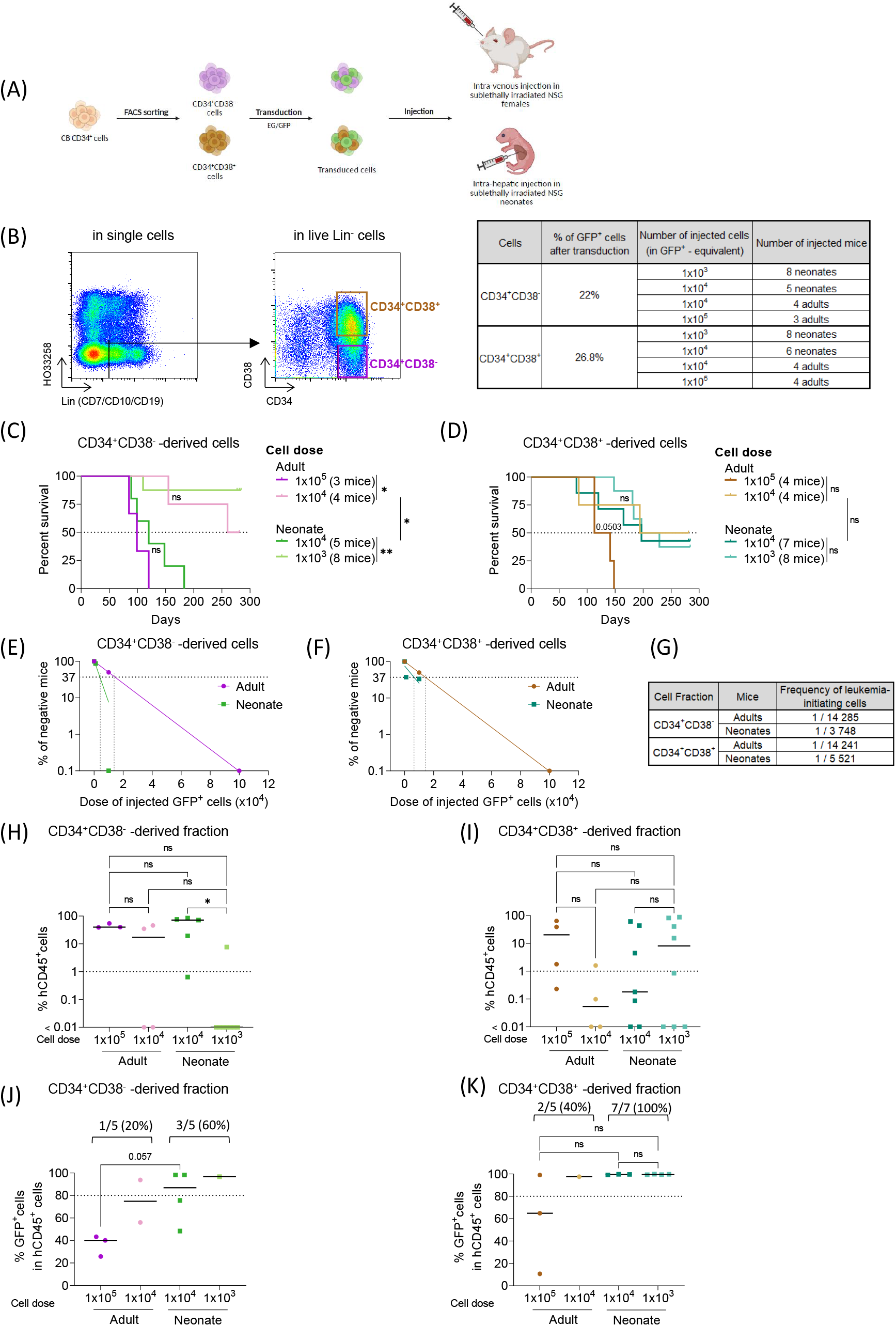
Both CD34^+^CD38^-^ and CD34^+^CD38^+^ cell fractions are sensitive to EG-driven transformation in neonate NSG mice. (A) Experimental design (*Created in BioRender. Gachet, S. (2026)* https://BioRender.com/0vvx4qo). CB CD34^+^CD38^-^ (immature compartment) and CD34^+^CD38^+^ (progenitor compartment) cells were flow-sorted, transduced and injected into sublethally irradiated (1Gy) NSG neonates (48h after birth by IH route, females and males, 1x10^3^ or 1x10^4^ GFP^+^ cells per mouse) or sublethally irradiated (2Gy) NSG female adults (IV route, 1x10^4^ or 1x10^5^ GFP^+^ cells per mouse). Neonates were weaned at the age of 28 days. Mice were monitored daily and euthanized when presenting disease symptoms (hind limb paralysis, signs of anemia, loss of weight). (B) Sorting strategy. Purified hematopoietic fractions were sorted according to CD34 and CD38 cell surface marker expression as indicated. Dot plots of the sorting strategy are shown (left panel). Table recapitulating the number of injected mice/condition. n=1 biological replicate done with CD34^+^CD38^-^ and CB CD34^+^CD38^+^ cells isolated from a pool of multiple CB samples. The transduction efficiency (% of GFP^+^ cells after transduction) is indicated (right panel). (C) Kaplan-Meier survival plot of NSG mice injected with EG-transduced CB CD34^+^CD38^-^ cells. Median survival was 99 days for adults receiving 1x10^5^ GFP^+^ cells, 270 days for adults receiving 1x10^4^ GFP^+^ cells and 120 days for neonates receiving 1x10^4^ GFP^+^ cells. (D) Kaplan-Meier survival plot of NSG mice injected with EG-transduced CB CD34^+^CD38^+^ cells. Median survival was 127 days for adults receiving 1x10^5^ GFP^+^ cells, 237 days for adults receiving 1x10^4^ GFP^+^ cells, 197 days for neonates receiving 1x10^4^ GFP^+^ cells and 213 days for neonates receiving 1x10^3^ GFP^+^ cells. (E-F) Frequency of leukemia-initiating cells in adults vs neonates. The fraction of non-responding (healthy) mice was plotted vs the number of injected CB CD34^+^CD38^-^ (E) or CB CD34^+^CD38^+^ (F) EG-GFP^+^ cells. The frequency of leukemia-initiating cells was estimated using the L-Calc software (StemCell Technologies). (G) Table summary of the frequency of leukemia-initiating cells in the CD34^+^CD38^-^ and CD34^+^CD38^+^ cell fractions in adults and neonates. (H-I) Percent of engrafted human CD45^+^ cells in the long bones BM of studied mice injected with CB CD34^+^CD38^-^ (H) and CD34^+^CD38^+^ (I) cells. (J-K) Percent of GFP^+^ cells in the long bones BM of studied mice (gated in hCD45^+^ cells) injected with CB CD34^+^CD38^-^ (J) and CD34^+^CD38^+^ (K) cells. Statistical significance is indicated as p values (C-D: Log-rank test, H-I-K: Kruskal-Wallis test, J: Mann-Whitney test). ns: not significant, *: p<0.05, **: p<0.01.

Overall, we uncovered that the intrinsic permissiveness of CB HSPCs to EG-mediated transformation is strongly influenced by the developmental stage of surrounding niche/microenvironment. Indeed, neonate NSG recipients were more supportive than adults to EG-driven leukemogenesis, making them a physiologically-relevant model for studying pediatric leukemogenesis with low cell inputs. While the recipient age does not influence the overall human cell engraftment dynamics with similar human cell levels being detected in neonate- and adult-derived mice, supportive extrinsic signals from the neonate environment are important determinants of the transformation. Now, it is not excluded that transplanting high cell numbers in a small body may also enhance autologous cytokine production favoring reciprocal cell cross-talks and potentially facilitate leukemia development, something we did not measure in this study. These growth factors may originate from normal (untransduced GFP^-^) and/or abnormal (EG/GFP^+^) cells and should be the focus of future investigations, as other factors than those identified previously^8^ may promote leukemia development from CB cells. Calculation of LIC activity unravelled higher (∼10-fold) frequencies in CD34^+^CD38^-/+^ cells than in total CD34^+^ cells in both adult and newborn recipients, highlighting heterogeneous CB cell subpopulations sensitive to EG-driven leukemogenesis. Therefore, identifying the rare transformation-permissive subpopulations within CB HSPCs as well as studying how EG fusion molecularly promotes transformation in the different subpopulations will also be of critical interest. Together, this novel model of setting EG-driven leukemogenesis in age-relevant recipients will be an interesting tool to dissect which and how extrinsic factors, including autocrine or paracrine cytokine production, facilitate leukemogenesis, and may provide relevant cell and molecular targets for therapeutic intervention.

## Author contributions

Klaudia Galant: formal analysis, investigation, methodology, project administration, validation, visualization, writing – original draft preparation, writing – review and editing Vilma Barroca: investigation, resources

Saiyirami Devanand: investigation, resources

Laurent Renou: investigation

Didier Busso: resources

Guillaume Piton: resources

Nathalie Dechamps: resources

Thomas Mercher: conceptualization, funding acquisition, methodology

Stéphanie Gachet: investigation, writing – original draft preparation, writing – review and editing

Françoise Pflumio: conceptualization, funding acquisition, methodology, project administration, resources, supervision, validation, writing – original draft preparation, writing – review and editing

## Acknowledgments

We thank the members of IRCM platforms for their help in the work. We acknowledge Julien Calvo for critical reading and editing the manuscript. We are undebted to T Domet and L Faivre from the Centre de Thérapie Cellulaire in Hôpital Saint Louis, Paris, France, for supplying cord blood samples.

This study was supported by funds from Inserm, CEA, University Paris Cité, University Paris Saclay, Institut National Du Cancer (PLBIO-2018-169, CONECT-AML: Subvention INCA-ARC-LIGUE_11905, PEDIAC: INCA_15670, PEDIAMOD22-008),

Ligue contre le cancer (FP and TM: équipes labellisées, KG: 4^th^ year of PhD funding TDHJ24888). FP and TM are members of the OPALE Carnot institute and of the Paris Kids Cancer program. FP is part of IHU La Leucémie Paris St Louis.

## Ethic statements

### Human sample collection

Umbilical cord blood (CB) samples were collected from healthy infants at the Cell Therapy Department in Hôpital Saint Louis, Paris, France with informed consent of the mothers based on the declaration of Helsinki. Samplings and experiments were acknowledged by the Institutional Review Board of INSERM (Opinion number 12–079, IRB00003888).

### Mouse experiments

All procedures were done in accordance with the recommendations of the European Community and French Ministry of Agriculture regulations (animal facility registration number: A920322) for the care and use of laboratory animals. Experimental procedures were specifically approved by the local Ethical Committee (CEEA 26: A18_105 and APAFIS #20538–2019050710555633).

NOD.Cg-PrkdcscidIl2rgtm1Wjl/SzJ (NSG) mice were originally obtained from the Jackson Laboratory (Bar Harbor, Maine, USA), housed and bred in specific pathogen-free animal facilities (IRCM, CEA, Fontenay-aux-Roses, France).

## References

1. Appelbaum FR, Gundacker H, Head DR, et al. Age and acute myeloid leukemia. Blood. 2006;107(9):3481–3485. doi:10.1182/blood-2005-09-3724

2. Blais S, Boutroux H, Pasquet M, et al. Is Acute Myeloblastic Leukemia in Children Under 2 Years of Age a Specific Entity? A Report from the FRENCH ELAM02 Study Group. Hemasphere. 2019;3(6):e316. doi:10.1097/HS9.0000000000000316

3. Hara Y, Shiba N, Yamato G, et al. Patients aged less than 3 years with acute myeloid leukaemia characterize a molecularly and clinically distinct subgroup. British Journal of Haematology. 2020;188(4):528–539. doi:10.1111/bjh.16203

4. de Rooij JDE, Branstetter C, Ma J, et al. Pediatric non–Down syndrome acute megakaryoblastic leukemia is characterized by distinct genomic subsets with varying outcomes. Nature Genetics. 2017;49(3):451–456. doi:10.1038/ng.3772

5. de Rooij JDE, Masetti R, van den Heuvel-Eibrink MM, et al. Recurrent abnormalities can be used for risk group stratification in pediatric AMKL: a retrospective intergroup study. Blood. 2016;127(26):3424–3430. doi:10.1182/blood-2016-01-695551

6. Hara Y, Shiba N, Ohki K, et al. Prognostic impact of specific molecular profiles in pediatric acute megakaryoblastic leukemia in non-Down syndrome. Genes, Chromosomes and Cancer. 2017;56(5):394–404. doi:10.1002/gcc.22444

7. Lopez CK, Noguera E, Stavropoulou V, et al. Ontogenic Changes in Hematopoietic Hierarchy Determine Pediatric Specificity and Disease Phenotype in Fusion Oncogene–Driven Myeloid Leukemia. Cancer Discov. 2019;9(12):1736–1753. doi:10.1158/2159-8290.CD-18-1463

8. Alonso-Pérez V, Galant K, Boudia F, et al. Developmental interplay between transcriptional alterations and a targetable cytokine signaling dependency in pediatric ETO2::GLIS2 leukemia. Mol Cancer. 2024;23(1):1–24. doi:10.1186/s12943-024-02110-y

9. Thiollier C, Lopez CK, Gerby B, et al. Characterization of novel genomic alterations and therapeutic approaches using acute megakaryoblastic leukemia xenograft models. Journal of Experimental Medicine. 2012;209(11):2017–2031. doi:10.1084/jem.20121343

10. Thirant C, Ignacimouttou C, Lopez CK, et al. ETO2-GLIS2 Hijacks Transcriptional Complexes to Drive Cellular Identity and Self-Renewal in Pediatric Acute Megakaryoblastic Leukemia. Cancer Cell. 2017;31(3):452–465. doi:10.1016/j.ccell.2017.02.006

11. Benbarche S, Lopez CK, Salataj E, et al. Screening of ETO2-GLIS2–induced Super Enhancers identifies targetable cooperative dependencies in acute megakaryoblastic leukemia. Science Advances. 2022;8(6):eabg9455. doi:10.1126/sciadv.abg9455

12. Le Q, Hadland B, Smith JL, et al. CBFA2T3-GLIS2 model of pediatric acute megakaryoblastic leukemia identifies FOLR1 as a CAR T cell target. J Clin Invest. 2022;132(22). doi:10.1172/JCI157101

13. Aid Z, Robert E, Lopez CK, et al. High caspase 3 and vulnerability to dual BCL2 family inhibition define ETO2::GLIS2 pediatric leukemia. Leukemia. 2023;37(3):571–579. doi:10.1038/s41375-022-01800-0

14. Gress V, Roussy M, Boulianne L, et al. CBFA2T3::GLIS2 Pediatric Acute Megakaryoblastic Leukemia is Sensitive to BCL-XL Inhibition by Navitoclax and DT2216. Blood Advances. Published online September 20, 2023:bloodadvances.2022008899. doi:10.1182/bloodadvances.2022008899

15. Boudia F, Baille M, Babin L, et al. Progressive chromatin rewiring by ETO2::GLIS2 revealed in a human iPSC model of pediatric leukemia initiation. Blood. Published online December 10, 2024:blood.2024024505. doi:10.1182/blood.2024024505

16. Lee GY, Jeong SY, Lee HR, Oh IH. Age-related differences in the bone marrow stem cell niche generate specialized microenvironments for the distinct regulation of normal hematopoietic and leukemia stem cells. Sci Rep. 2019;9(1):1007. doi:10.1038/s41598-018-36999-5

17. Takihara Y, Higaki T, Yokomizo T, et al. Bone marrow imaging reveals the migration dynamics of neonatal hematopoietic stem cells. Commun Biol. 2022;5(1):776. doi:10.1038/s42003-022-03733-x

18. Sánchez-Lanzas R, Jiménez-Pompa A, Ganuza M. The evolving hematopoietic niche during development. Front Mol Biosci. 2024;11. doi:10.3389/fmolb.2024.1488199

19. Ishikawa F, Yoshida S, Saito Y, et al. Chemotherapy-resistant human AML stem cells home to and engraft within the bone-marrow endosteal region. Nat Biotechnol. 2007;25(11):1315–1321. doi:10.1038/nbt1350

20. Dykstra C, Lee AJ, Lusty EJ, et al. Reconstitution of immune cell in liver and lymph node of adult- and newborn-engrafted humanized mice. BMC Immunology. 2016;17(1):18. doi:10.1186/s12865-016-0157-9

21. Keng CT, Sze CW, Zheng D, et al. Characterisation of liver pathogenesis, human immune responses and drug testing in a humanised mouse model of HCV infection. Gut. 2016;65(10):1744–1753. doi:10.1136/gutjnl-2014-307856

22. Her Z, Yong KSM, Paramasivam K, et al. An improved pre-clinical patient-derived liquid xenograft mouse model for acute myeloid leukemia. J Hematol Oncol. 2017;10(1):1–14. doi:10.1186/s13045-017-0532-x

23. Sippel TR, Radtke S, Olsen TM, Kiem HP, Rongvaux A. Human hematopoietic stem cell maintenance and myeloid cell development in next-generation humanized mouse models. Blood Adv. 2019;3(3):268–274. doi:10.1182/bloodadvances.2018023887

24. Beyer AI, Muench MO. Comparison of Human Hematopoietic Reconstitution in Different Strains of Immunodeficient Mice. Stem Cells Dev. 2017;26(2):102–112. doi:10.1089/scd.2016.0083

25. Notta F, Doulatov S, Dick JE. Engraftment of human hematopoietic stem cells is more efficient in female NOD/SCID/IL-2Rgc-null recipients. Blood. 2010;115(18):3704–3707. doi:10.1182/blood-2009-10-249326

26. Mian SA, Ariza-McNaughton L, Anjos-Afonso F, et al. Influence of donor– recipient sex on engraftment of normal and leukemia stem cells in xenotransplantation. Hemasphere. 2024;8(5):e80. doi:10.1002/hem3.80

27. Doulatov S, Notta F, Laurenti E, Dick JE. Hematopoiesis: A Human Perspective. Cell Stem Cell. 2012;10(2):120–136. doi:10.1016/j.stem.2012.01.006

28. Karamitros D, Stoilova B, Aboukhalil Z, et al. Single-cell analysis reveals the continuum of human lympho-myeloid progenitor cells. Nat Immunol. 2018;19(1):85–97. doi:10.1038/s41590-017-0001-2

