## Supplemental materials and figures for "Transplantation in neonate mouse recipients enhances umbilical cord blood CD34^+^ cells permissiveness to ETO2::GLIS2-driven transformation"

**Supplemental files**

#### Material and Methods

##### Human sample collection

Umbilical cord blood (CB) samples were collected from healthy infants at Cell Therapy Department in Hôpital Saint Louis, Paris, France with informed consent of the mothers based on the declaration of Helsinki. Samplings and experiments were acknowledged by the Institutional Review Board of INSERM (Opinion number 12-079, IRB00003888, FWA00005831).

##### $CD34^+$ cell isolation

CB cells were subjected to Ficoll gradient (Lymphocyte Separation Medium, Eurobio Scientific) and enriched for $CD34^+$ cells using the human CD34 MicroBeads Kit according to manufacturer instructions (Miltenyi Biotec). Purity (>60%) of $CD34^+$ cell suspensions was tested using immunostaining (anti-CD34 antibody, clone 581, BioLegend). Cells were used fresh or after freezing/thawing process in fetal bovine serum (FBS, Sigma) supplemented with 10% DMSO (Sigma). Only samples with ≥70% viable cells were used.

##### Hematopoietic subfractions cell sorting

One day prior to sorting, CB $CD34^+$ cells were thawed in a 37°C bath and resuspended in complete medium (IMDM + Glutamax (Gibco) supplemented with 20% of FBS and 1% P/S (Gibco)). Cells were spun at 300g for 5min to remove the DMSO, resuspended in the same medium and kept overnight in presence of DNAse I (10 µg/mL, Roche). Cells were stained with a cocktail of antibodies (Supplemental Table 1) for 15-20min at 4°C, then washed with DPBS. Before cell sorting, cells were spun down at 300g for 5min and resuspended in DPBS (Gibco) at 25-30x$10^6$ cells/mL. Cells were sorted on a BD Influx™ Cell Sorter (BD Biosciences). Compensation controls were performed using single-stained compensation beads (UltraComp eBeads™ Plus Compensation Beads, Invitrogen). Dead cells were excluded using Hoechst 33258 dye (Invitrogen).

##### Lentiviral vector production and transduction

Concentrated lentiviral supernatants were generated and produced at the Genetic engineering and protein biochemistry (CiGex) platform of CEA/DRCM. The lentiviral pTrip-MND-EG/eGFP construct was used to express the *EG* fusion.

Purified $CD34^+$ and sorted $CD34^+CD38^{+/-}$ cells (1x$10^6$ cells/mL) were seeded in BIT 9500 Serum Substitute (StemCell Technologies) supplemented with 1% P/S, 100 ng/mL hSCF, 100 ng/mL hFLT3L, 60 ng/mL hIL3 and 200 ng/mL hTPO (all from Miltenyi Biotec). Transduction lasted for 2 to 3 days at 37°C 5% $CO_2$ in presence of *EG* vectors (MOI=10). Transduced cells were washed three times using DPBS before further use. Transduction efficiency (% of $GFP^+$ cells) was analyzed on the BD FACSCanto™ II analyzer (BD Biosciences). Cells were injected after 2 days of culture (1x$10^5$ cells/mL, in IMDM + Glutamax, 15% FBS, 1% P/S, 100 ng/mL hSCF, 100 ng/mL hFLT3L, 60 ng/mL hIL3, 200 ng/mL hTPO, 2 U/mL of hEPO (PeproTech)) when performing xenotransplantation assays.

**Xenotransplantation**

All procedures were done in accordance with the recommendations of the European Community and French Ministry of Agriculture regulations (animal facility agreement number: E92-032-02, delivered 26 May 2024) for the care and use of laboratory animals. Experimental procedures were specifically approved by the local Ethical Committee (CEEA 26: A18_105 and APAFIS #20538-2019050710555633). NOD.Cg-PrkdcscidIl2rgtm1Wjl/SzJ (NSG) were originally obtained from the Jackson Laboratory (Bar Harbor, Maine, USA), housed and bred in specific pathogen-free animal facilities (IRCM, CEA, Fontenay-aux-Roses, France).

NSG neonates were obtained after timed matings of 12- to 24-week-old mice. Pregnancy was confirmed the next morning by observation of a vaginal plug. Neonates were sublethally irradiated at 1 Gy 48 hours after birth, together with the dam to avoid rejection by the mother. Cell transplant was performed 1-2 hours after irradiation by intrahepatic (or by intravenous, one experiment) route. Briefly, neonates were gently restrained, snout facing upwards. A 30G insulin syringe loaded with 0.1-1x$10^5$ $GFP^+$-transduced cells/50µL was inserted in the liver. For intravenous neonate injection, 0.1-1x$10^5$ $GFP^+$-transduced cells/50µL were injected directly in the facial vein using a 30G insulin syringe. After injection, neonates were moved back to the cage with the dam, and monitored closely for 48-72 hours for survival or rejection by the mother. Both female and male neonates were used in these experiments. Neonates were weaned 28 days after birth.

8 to 12-week-old adult NSG female mice were sublethally irradiated at 2 Gy, before intravenous retro-orbital cell transplant under isoflurane anesthesia. Mice received 0.1-1x$10^6$ $GFP^+$-transduced cells/100µL. In some experiments, mice were injected by intra-bone route (20µL) directly in the femur after buprenorphine analgesia and under isoflurane anesthesia.

Mice were monitored daily and euthanized when exhibiting disease symptoms (hind limbs paralysis, anemia, loss of weight) or >8 months after transplant if not sick to explore human hematopoietic cell development. Tibiae/femora (4 long bones, BM) were harvested and hematopoietic cells were recovered after flushing BM with DPBS using a 23G gauge needle. The isolated cells were kept at 4°C on ice during the process. The frequency of leukemia-initiating cells was estimated using the L-Calc software (StemCell Technologies) by plotting the percent of healthy mice (mice without leukemia) against the number of injected $GFP^+$ cells.

**Flow cytometry**

Cells were stained with human specific antibodies purchased from BD Biosciences, BioLegend or eBioscience (Supplemental Table 2), in DPBS for 15-30min at 4°C. Labelled cells were analyzed on a BD FACSCanto™ II or BD LSR™ II using the FACS Diva software (BD Biosciences). Compensation controls were performed using single-stained compensation beads (UltraComp eBeads™ Plus Compensation Beads,

Invitrogen). After acquisition, live cell analysis was done using FlowJo™ v10, excluding debris and doublets using forward and light scatter and dead cells by exclusion of the Zombie Aqua™ Viability Dye (1:500, BioLegend).

**Statistical analysis**

Statistical analyses were performed with GraphPad Prism 9.4. Data were analyzed by Mann-Whitney or Kruskal-Wallis tests followed by a multiple comparison test (as indicated in figures).

For xenotransplantation analysis, Kaplan–Meier survival curves were performed and the statistical significance was determined with the Log-rank test. Overall survival was defined as the time from cell injection to euthanasia or death at predefined endpoints. Animals were censored if the cause of death/sacrifice was not leukemia related.

Data are presented as medians for *in vivo* experiments. Results were considered statistically significant if $p<0.05$. p values are indicated by "ns" when not significant, * when $p \leq 0.05$, ** when $p \leq 0.01$, *** when $p \leq 0.001$ and **** when $p \leq 0.0001$.

#### Supplemental Tables

##### Supplemental Table 1: antibodies used for flow-sorting CD34$^+$CD38$^-$ and CD34$^+$CD38$^+$ hematopoietic populations

| Antibody | Fluorochrome | Clone | Supplier | Dilution |
|---|---|---|---|---|
| CD7 | FITC | CD7-6B7 | BioLegend | 1:100 |
| CD10 | FITC | SN5c | BioLegend | 1:100 |
| CD19 | FITC | HIB1 | BioLegend | 1:100 |
| CD34 | PerCPCy5.5 | 581 | BioLegend | 1:100 |
| CD38 | APC-eFluor 780 | HIT2 | eBiosciences | 1:100 |
| HO33258 | Viability | / | Invitrogen | / |

##### Supplemental Table 2: antibodies used for analysis of cells recovered from deceased recipient mice

| Antibody | Fluorochrome | Clone | Supplier | Dilution |
|---|---|---|---|---|
| CD3 | PerCPCy5.5 | SK7 | BioLegend | 1:100 |
| CD11b | PECy7 | ICRF44 | BioLegend | 1:100 |
| CD19 | PE | HIB19 | BioLegend | 1:100 |
| CD33 | PE | P67.6 | BioLegend | 1:100 |
| CD34 | PECy7 | 581 | BioLegend | 1:100 |
| CD41 | PerCPCy5.5 | HIP8 | BioLegend | 1:100 |
| CD45 | APC-eFluor780 | HI30 | eBiosciences | 1:50 |
| CD56 | APC | CMSSB | eBiosciences | 1:100 |
| KIT/CD117 | BV421 | 104D2 | BioLegend | 1:50 |
| Zombie Aqua | Viability | / | BioLegend | 1:500 |

### Figure S1

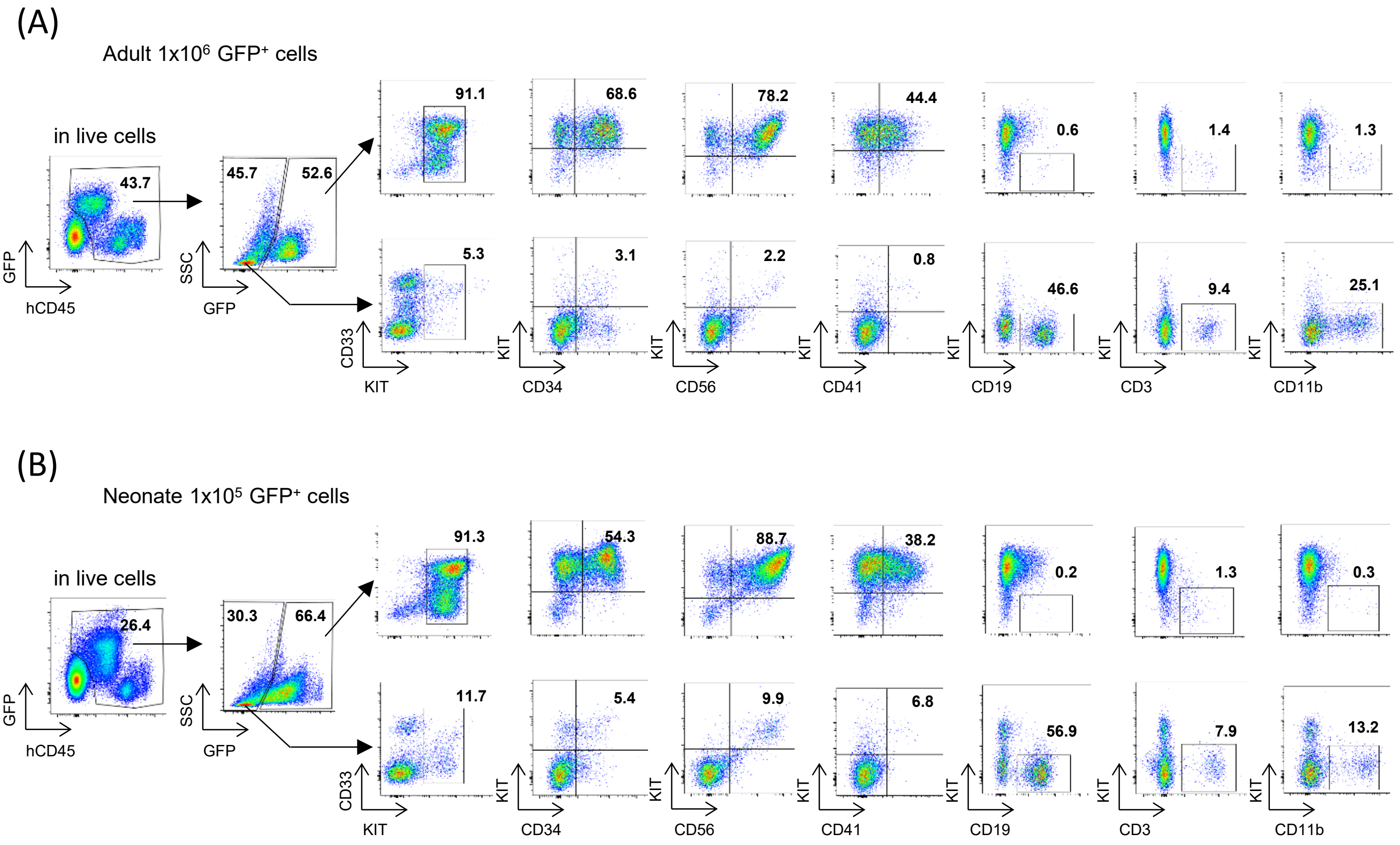


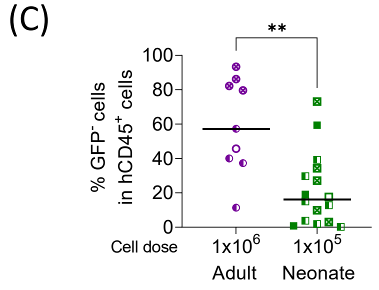


**Figure S1: Gating strategy employed for human cell analysis in deceased mice.** (A) Representative phenotype analysis of the human hematopoietic cells recovered from long bones BM of an adult deceased recipient injected with $1x10^6$ $GFP^+$ cells. (B) Representative phenotype analysis of the human hematopoietic cells recovered from long bones BM of a neonate deceased recipient injected with $1x10^5$ $GFP^+$ cells. Both mice in (A) and (B) are from the same experiment, and got sick 174 days and 153 days after cell inoculation, respectively. (C) Percent of $GFP^-$ cells in the long bones BM of deceased recipients (gated in $hCD45^+$ cells). Symbols are used as in Fig1D-G and show the different experiments depicted in Fig1A. Statistical significance is indicated as p values (C: Mann-Whitney test). **: $p<0.01$.

### Figure S2

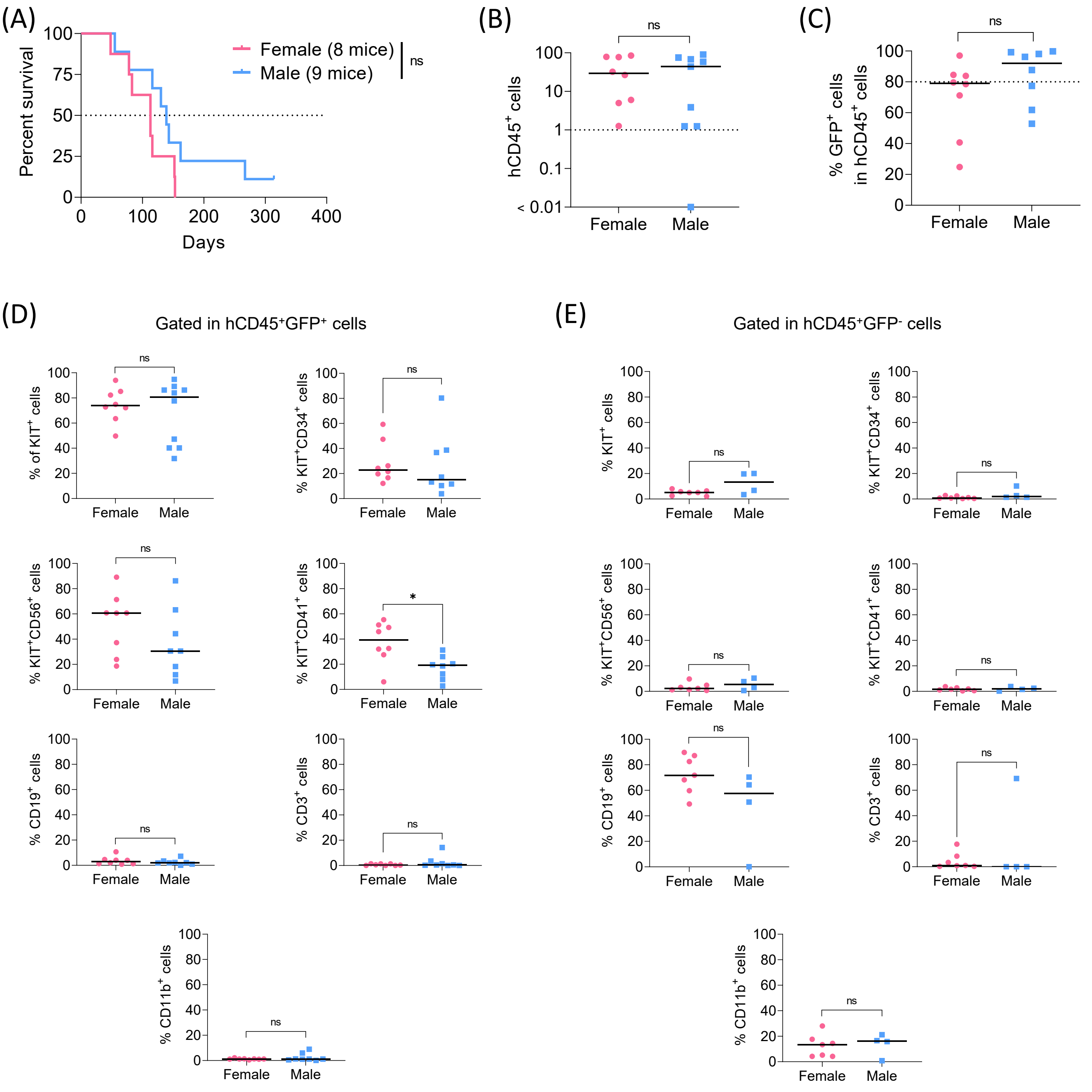


**Figure S2: Repopulation by the transplanted CB EG cells does not differ between female and male NSG neonates.** (A) Kaplan-Meier survival plot of NSG neonates injected with 1x$10^5$ EG-transduced CB cells. Median survival was 113 days for female neonates, 139 days for male neonates. (B) Percent of engrafted human $CD45^+$ cells in the long bones BM of deceased recipients. (C) Percent of $GFP^+$ cells in the long bones BM of deceased recipients (gated in $hCD45^+$ cells). (D) Percent of $KIT^+$, $KIT^+CD34^+$, $KIT^+CD56^+$, $KIT^+CD41^+$ and normal $CD19^+$, $CD3^+$ and $CD11b^+$ cells in the long bones BM of deceased recipients (gated in $hCD45^+GFP^+$ cells). (E) Percent of $KIT^+$, $KIT^+CD34^+$, $KIT^+CD56^+$, $KIT^+CD41^+$ and normal $CD19^+$, $CD3^+$ and $CD11b^+$ cells in the long bones BM of deceased recipients (gated in $hCD45^+GFP^-$ cells). Statistical significance is indicated as p values (A: Log-rank test, B-C-D-E: Mann-Whitney test). ns: not significant, *: $p<0.05$.

#### Figure S3

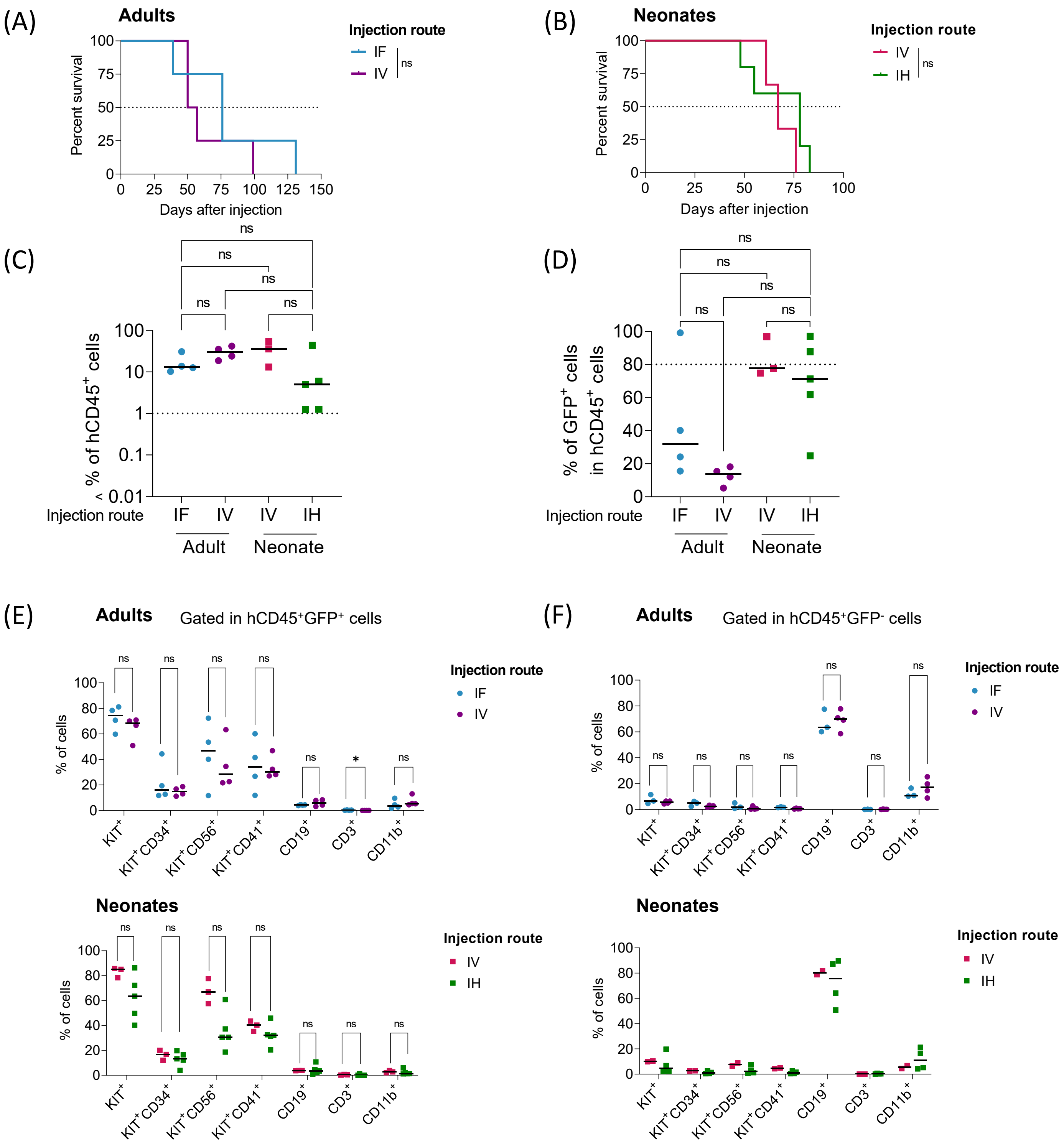
(A)
Adults
Percent survival
Days after injection
Injection route
IF
IV
ns
(B)
Neonates
Percent survival
Days after injection
Injection route
IV
IH
ns
(C)
% of hCD45+ cells
Injection route
IF
IV
IV
IH
Adult
Neonate
(D)
% of GFP+ cells in hCD45+ cells
Injection route
IF
IV
IV
IH
Adult
Neonate
(E)
Adults
Gated in hCD45+GFP+ cells
% of cells
Injection route
IF
IV
KIT+
KIT+CD34+
KIT+CD56+
KIT+CD41+
CD19+
CD3+
CD11b+
Neonates
Injection route
IV
IH
(F)
Adults
Gated in hCD45+GFP- cells
Injection route
IF
IV
Neonates
Injection route
IV
IH

**Figure S3: Survival and repopulation of adult and neonate recipients is independent of the injection route.** (A) Kaplan-Meier survival plot of NSG adult mice injected with $1x10^6$ EG-transduced CB cells by intra-femoral route (IF, 4 mice) or intravenously (IV, 4 mice). Median survival was 76 days for mice injected by IF route, 53.5 days for mice injected by IV route. (B) Kaplan-Meier survival plot of NSG neonates injected with $1x10^5$ EG-transduced CB cells intravenously (IV, 3 mice) or by intrahepatic route (IH, 5 mice). Median survival was 67 days for mice injected by IV route, 78 days for mice injected by IH route. (C) Percent of engrafted human $CD45^+$ cells in the long bones BM of deceased recipients. (D) Percent of $GFP^+$ cells in the long bones BM of deceased recipients (gated in $hCD45^+$ cells). (E) Percent of $KIT^+$, $KIT^+CD34^+$, $KIT^+CD56^+$, $KIT^+CD41^+$ and normal $CD19^+$, $CD3^+$ and $CD11b^+$ cells in the long bones BM of deceased recipients (gated in $hCD45^+GFP^+$ cells). Upper panel: adult recipients. Lower panel: neonate recipients. (F) Percent of $KIT^+$, $KIT^+CD34^+$, $KIT^+CD56^+$, $KIT^+CD41^+$ and normal $CD19^+$, $CD3^+$ and $CD11b^+$ cells in the long bones BM of deceased recipients (gated in $hCD45^+GFP^-$ cells). Upper panel: adult recipients. Lower panel: neonate recipients. Statistical significance is indicated as p values (A-B: Log-rank test, C-D: Kruskal-Wallis test, E-F: Mann-Whitney test). ns: not significant, *: $p<0.05$.

### Figure S4

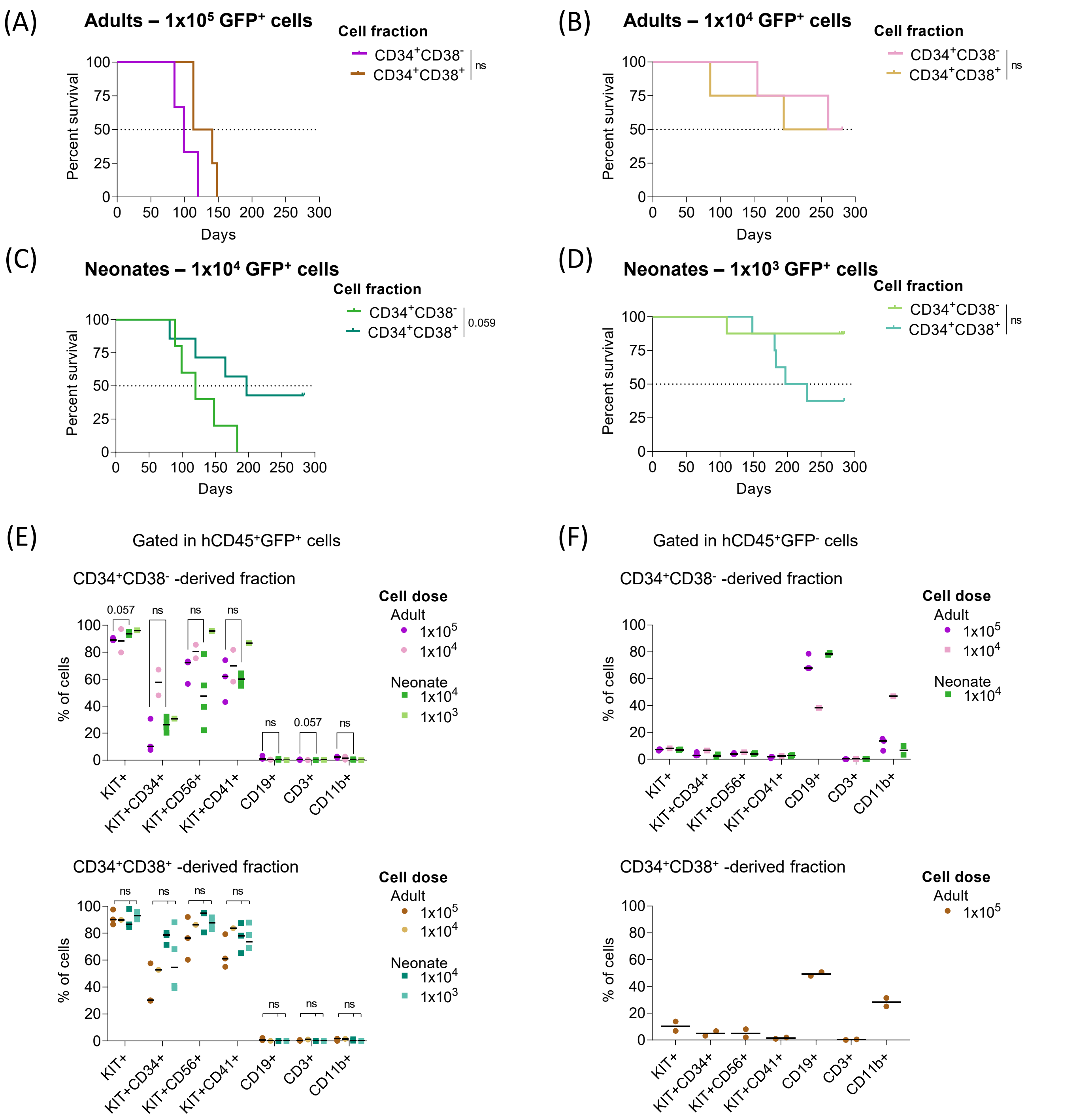

(A)
Adults – 1x10⁵ GFP⁺ cells
Cell fraction
CD34⁺CD38⁻
CD34⁺CD38⁺
ns
Percent survival
Days
(B)
Adults – 1x10⁴ GFP⁺ cells
Cell fraction
CD34⁺CD38⁻
CD34⁺CD38⁺
ns
(C)
Neonates – 1x10⁴ GFP⁺ cells
Cell fraction
CD34⁺CD38⁻
CD34⁺CD38⁺
0.059
(D)
Neonates – 1x10³ GFP⁺ cells
Cell fraction
CD34⁺CD38⁻
CD34⁺CD38⁺
ns
(E)
Gated in hCD45⁺GFP⁺ cells
CD34⁺CD38⁻ -derived fraction
CD34⁺CD38⁺ -derived fraction
% of cells
Cell dose
Adult
1x10⁵
1x10⁴
Neonate
1x10⁴
1x10³
0.057
ns
KIT+
KIT+CD34+
KIT+CD56+
KIT+CD41+
CD19+
CD3+
CD11b+
(F)
Gated in hCD45⁺GFP⁻ cells
CD34⁺CD38⁻ -derived fraction
CD34⁺CD38⁺ -derived fraction

**Figure S4: Both $CD34^+CD38^-$ and $CD34^+CD38^+$ hematopoietic fractions are sensitive to EG transformation in a neonate NSG mouse model.** (A) Kaplan-Meier survival plot of NSG adult mice injected by IV route with $1x10^5$ EG-transduced $CD34^+CD38^-$ (3 mice) or $CD34^+CD38^+$ (4 mice) cells. Median survival was 99 days for mice injected with $CD34^+CD38^-$ cells, 127 days for mice injected with $CD34^+CD38^+$ cells. (B) Kaplan-Meier survival plot of NSG adult mice injected by IV route with $1x10^4$ EG-transduced $CD34^+CD38^-$ (4 mice) or $CD34^+CD38^+$ (4 mice) cells. Median survival was 270.5 days for mice injected with $CD34^+CD38^-$ cells, 237.5 days for mice injected with $CD34^+CD38^+$ cells. (C) Kaplan-Meier survival plot of NSG neonate mice injected by IH route with $1x10^4$ EG-transduced $CD34^+CD38^-$ (5 mice) or $CD34^+CD38^+$ (6 mice) cells. Median survival was 120 days for mice injected with $CD34^+CD38^-$ cells, 197 days for mice injected with $CD34^+CD38^+$ cells. (D) Kaplan-Meier survival plot of NSG adult mice injected by IH route with $1x10^3$ EG-transduced $CD34^+CD38^-$ (8 mice) or $CD34^+CD38^+$ (8 mice) cells. Median survival was 213 days for mice injected with $CD34^+CD38^+$ cells. (E) Percent of $KIT^+$, $KIT^+CD34^+$, $KIT^+CD56^+$, $KIT^+CD41^+$ and normal $CD19^+$, $CD3^+$ and $CD11b^+$ cells in the long bones BM of deceased recipients (gated in $hCD45^+GFP^+$ cells). Upper panel: recipients injected with $CD34^+CD38^-$ cells. Lower panel: recipients injected with $CD34^+CD38^+$ cells. (F) Percent of $KIT^+$, $KIT^+CD34^+$, $KIT^+CD56^+$, $KIT^+CD41^+$ and normal $CD19^+$, $CD3^+$ and $CD11b^+$ cells in the long bones BM of deceased recipients (gated in $hCD45^+GFP^-$ cells). Upper panel: recipients injected with $CD34^+CD38^-$ cells. Lower panel: recipients injected with $CD34^+CD38^+$ cells. Statistical significance is indicated as p values (A-B-C-D: Log-rank test, E upper panel: Mann-Whitney test, E lower panel: Kruskal-Wallis test). ns: not significant.
